# Cellular and molecular signatures of *in vivo* GABAergic neurotransmission in the human brain

**DOI:** 10.1101/2021.06.17.448812

**Authors:** PB Lukow, D Martins, M Veronese, AC Vernon, P McGuire, FE Turkheimer, G Modinos

## Abstract

Diverse GABAergic interneuron microcircuits orchestrate information processing in the brain. Understanding the cellular and molecular composition of these microcircuits, and whether these can be imaged by available non-invasive *in vivo* methods is crucial for the study of GABAergic neurotransmission in health and disease. Here, we use human gene expression data and state-of-the-art imaging transcriptomics to uncover co-expression patterns between GABA_A_ receptor subunits and interneuron subtype-specific markers, and to decode the cellular and molecular signatures of gold-standard GABA PET radiotracers, [^11^C]Ro15-4513 and [^11^C]flumazenil. We find that the interneuron marker somatostatin is co-expressed with GABA_A_ receptor-subunit genes *GABRA5* and *GABRA2*, and their distribution maps onto [^11^C]Ro15-4513 binding *in vivo*. In contrast, the interneuron marker parvalbumin co-expressed with more predominant GABA_A_ receptor subunits (*GABRA1, GABRB2* and *GABRG2*), and their distribution tracks [^11^C]flumazenil binding *in vivo*. These results have important implications for the non-invasive study of GABAergic microcircuit dysfunction in psychiatric conditions.

## Introduction

Although accounting for less than 30% of cortical cells, GABAergic inhibitory interneurons control information processing throughout the brain (1–4). Their diverse functions include input gating into cortical (5) and subcortical (6) structures, regulating critical period boundaries, homeostasis (7), regulating local network activity and entraining cortical network oscillations (1). Due to their critical role in such wide range of brain functions, GABAergic interneuron dysfunction has been implicated in several psychiatric and neurological conditions, including affective disorders (6,8,9) and schizophrenia (7,10,11).

The GABAergic system comprises diverse interneuron subtypes, innervating different neural targets through a variety of receptors (1). This complexity poses a challenge to microcircuit investigation in the human brain *in vivo* that can be both selective and non-invasive. GABAergic interneurons vary in their firing threshold, spiking frequency and location of postsynaptic cell innervation, which makes them fit for various functions including the control of synaptic input into the local network and neuronal output regulation (5,12). This multitude of interneurons can be classified through the expression of specific proteins (interneuron markers) (13). While the vast majority of interneurons in the brain are positive for either parvalbumin (PVALB), somatostatin (SST), or vasoactive intestinal peptide (VIP), further specific subtypes can be identified through the expression of other markers, such as cholecystokinin (CCK) (3). This array of interneurons achieves fine-tuned inhibitory responses via the ionotropic GABA_A_ receptor which mediates postsynaptic cell hyperpolarisation. The GABA_A_ receptor (or GABA_A_R) is a pentameric chloride channel which most commonly comprises two α, two β and one γ subunit (14). There are five subtypes of the α subunit and three each of the β and γ subunits; moreover, β can be replaced by a θ subunit, and γ can be replaced by δ, ε or π (14). This generates a great variety of receptors, the biology and pharmacology of which are determined by their subunit composition. For instance, the low-affinity α1 subunit-containing GABA_A_R (GABA_A_Rα1) mediates phasic, or activity-dependent inhibition on the postsynaptic cell, whereas GABA_A_Rα5 present higher affinity to GABA, maintaining a more continuous inhibitory tone extrasynaptically (15–19).

Detailed investigation of the roles that GABAergic interneuron neurotransmission may play in brain function in health and disease requires: 1) precise knowledge of the basic principles underlying the organization of this complex system in the human brain; and 2) the ability to identify and tease apart its specific microcircuits. In this context, positron emission tomography (PET) allows *in vivo* GABA_A_R quantification in anatomically defined brain regions through the use of radiolabelled ligands, mainly [^11^C]Ro15-4513, with high affinity to GABA_A_Rα5, and [^11^C]flumazenil, a benzodiazepine site-specific ligand with affinity to GABA_A_Rα1-3 and α5 (20,21). Although receptor affinity for these radiotracers has been confirmed in preclinical research (22), their specificity for defined cell types is unknown. This compromises the understanding of which specific interneuron microcircuits contribute the most to inter-regional differences in signals obtained from human GABA PET measurements. Interestingly, both the distribution of GABAergic interneurons and GABA PET radiotracer binding are heterogeneous across the brain. For instance, *SST* and *PVALB* follow an anticorrelated distribution (11), as do the binding patterns of [^11^C]Ro15-4513 and [^11^C]flumazenil (23). Moreover, postsynaptic expression of GABA_A_R subunits, encoded by individual genes in target neurons, is associated with specific interneuron subtypes in rodent (19,24–26). Brain-wide gene expression atlases such as the Allen Human Brain Atlas (AHBA) are increasingly being used to gain insight into the mechanisms linking complex brain microcircuits to measurements of human brain function *in vivo* (27). Thus, determining the spatial relationships between the expression of GABA_A_R subunits and interneuron markers may inform the basic principles that govern the spatial organization major GABAergic interneuron microcircuits in the human brain. Moreover, because these brain-wide gene expression data can be integrated with neuroimaging measures, such as binding from GABA PET tracers, this approach may help understand which interneuron microcircuits follow and contribute the most to the spatial pattern of binding of these tracers.

Here, we used state-of-the-art imaging transcriptomics to uncover patterns of co-expression between GABA_A_R subunits and interneuron markers in the human brain, and to decode the molecular and cellular signatures of two gold-standard GABA PET radiotracers, [^11^C]Ro15-4513 and [^11^C]flumazenil (28). We demonstrate that *SST* co-expresses with two GABA_A_R subunits implicated in affective function, *GABRA5* and *GABRA2*, while *PVALB* strongly correlates with genes encoding subunits of the most prevalent GABAergic receptor in the brain, GABA_A_Rα1β2γ2. While [^11^C]Ro15-4513 signal covaries with the expression of *SST*, *GABRA5, GABRA2* and *GABRA3*, [^11^C]flumazenil signal is positively correlated with the expression of *PVALB* and genes of the GABA_A_Rα1β2γ2 receptor. We also show that *VIP* is co-expressed with *CCK*, and that these two genes covary with both radiotracers. Taken together, our findings show for the first time in human that 1) the expression of markers for PVALB and SST interneurons is associated with distinct GABA_A_ receptor complexes; and 2) that the distribution of genes from those two interneuron populations can be tracked by [^11^C]Ro15-4513 and [^11^C]flumazenil binding *in vivo*. Given the key role for PVALB and SST cells in healthy brain function, and the strong implication of PVALB and SST dysfunction in several brain disorders (8), our findings provide a detailed framework to inform future GABA PET studies in the choice of PET tracer to investigate specific GABAergic microcircuitry in health and disease.

## Results

### GABAergic interneuron markers co-express with specific GABA_A_R subunits

Our first aim was to identify co-expression patterns between interneuron cell-type markers and GABA_A_R subunits prior to their integration with the PET imaging data. Interneuron markers of interest comprised: the GABA-synthesising enzymes GAD67 (*GAD1*) and GAD65 (*GAD2*) (29), parvalbumin (*PVALB*) (25), somatostatin (*SST*) (30), vasoactive intestinal peptide (*VIP*) (31), cholecystokinin (*CCK*),neuropeptide Y (*NPY*) (32), calbindin (*CALB1*) (30), calretinin (*CALB2*) (30), neuronal nitric oxide synthase (*NOS1*) (3), reelin (*RELN*) (33), and the tachykinin precursor genes *TAC1, TAC3* and *TAC4* (34), selected according to preclinical literature and Petilla classification of GABAergic interneurons (13). Data on all available GABA_A_R subunits passing quality threshold were included: (α1-5 (*GABRA1-5*), β1-3 (*GABRB1-3*), γ1-3 (*GABRG1-3*), ε (*GABRE*) and δ (*GABRD*)). Thus, the subunits α6 (*GABRA6*), π (*GABRP*), θ (*GABRQ*) and ρ1-3 (*GABRR1-3*) were not used in further analyses as they did not show levels of expression above background. We performed weighted gene co-expression network analysis (WGCNA) (35) on gene expression data from the AHBA (36). This dataset contained microarray data on 15,633 genes from six *post-mortem* samples across the left (n=6 healthy donors) and right hemispheres (n=2 healthy donors), which were resampled into 83 brain regions of the Desikan-Killiany atlas (37). WGCNA is a data-driven approach that allows to identify clusters (modules) of highly correlated genes across the whole transcriptome (35).

WGCNA identified 52 co-expression clusters, 13 of which included genes encoding interneuron markers and GABA_A_R subunits of interest. We selected these 13 clusters to investigate which genes shared cluster allocation (Figure 1A). *SST* was located in the same cluster as *GABRA5, GABRA2* and *GABRB1. PVALB* had its own cluster (i.e., it was not located in the same cluster as any other gene of interest). *VIP* was found in the same cluster as *CCK* and no other genes of interest. As those three individual clusters included three of the main non-overlapping interneuron markers, labelling the majority of GABAergic inhibitory cells in the mammalian brain (2), we investigated their enrichment in genes co-expressed in specific cell types defined by previous single-cell transcriptomic analysis (38) using the WEB-based GEne SeT AnaLysis Toolkit (39). These analyses revealed inhibitory interneuron cell-type enrichment in the *SST* and *PVALB* clusters, and excitatory cell-type enrichment in the *VIP* cluster (for full results, see Supplement). This indicated that the subsequent analyses including the *SST* and *PVALB* clusters were related to GABAergic interneuron cell-types. Other separate clusters of genes included *GAD1, GABRA1, GABRB2, GABRG2* and *GABRG3; CALB1, CALB2, GABRG1* and *GABRE; GABRA4, NPY* and *TAC3;* and *GAD2* and *NOS1*. Finally, *GABRB1, GABRB3, GABRA3, RELN, TAC1, GABRD* and *TAC4* were all found individually in separate clusters that did not share assignment with any other gene of interest.

**Figure 1.**
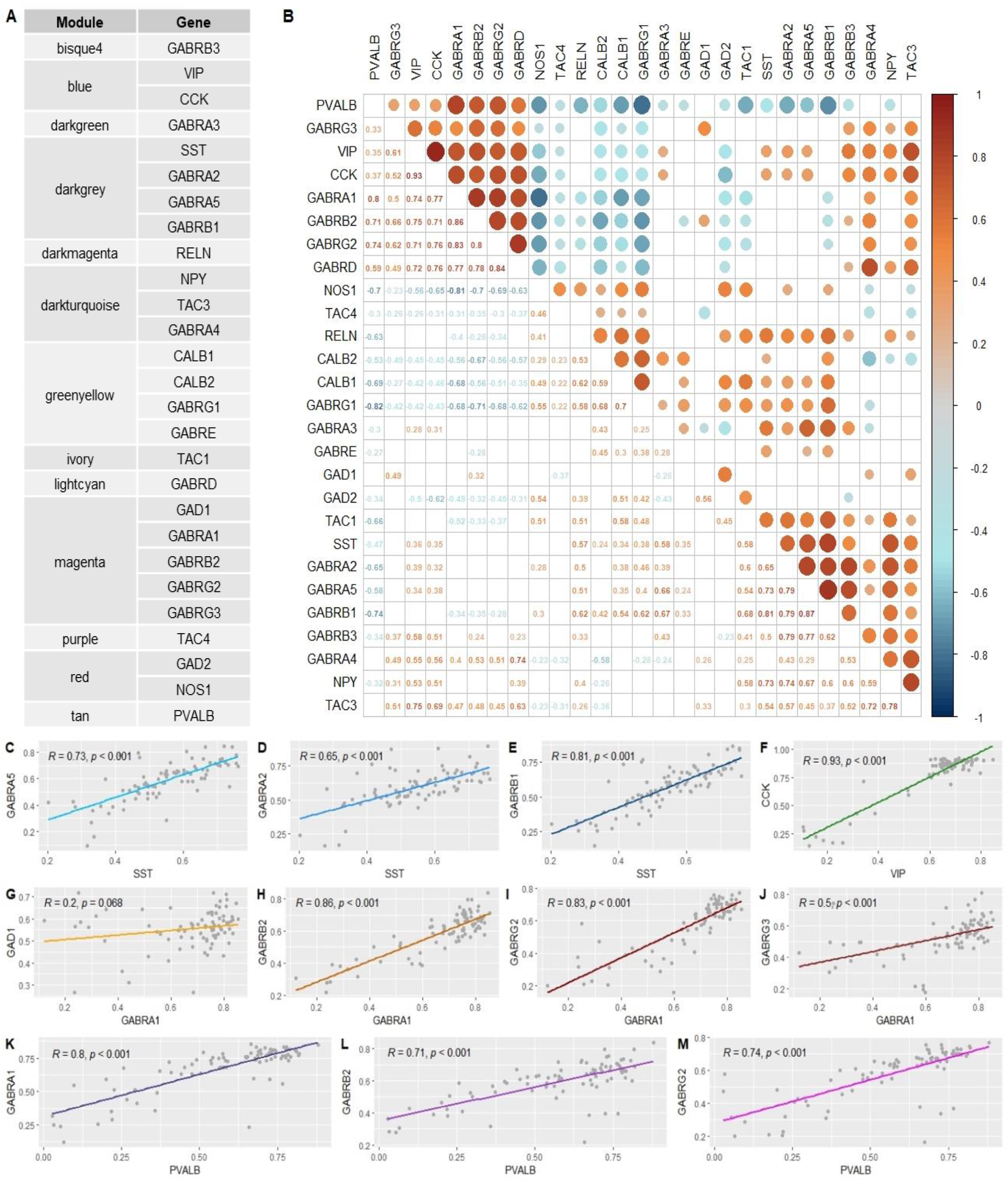
Specific GABAergic interneuron markers co-express with different GABA_A_R subunits. (**A**) Co-expression cluster assignment and (**B**) bivariate correlations (p<0.05) between GABAergic interneuron markers and GABA_A_R subunits. Pairwise correlations between (**C-E**) somatostatin (*SST*), (**F**) vasoactive intestinal peptide (*VIP*), (**G-J**) *GABRA1* and (**K-M**) parvalbumin (*PVALB*), and other genes of interest sharing their cluster assignment in the AHBA dataset. *GAD1*, GABA-synthesising enzyme GAD67, *GAD2*, GABA-synthesising enzyme GAD65 (*GAD2*), *CCK*, cholecystokinin, *NPY*, neuropeptide Y, *CALB1*, calbindin, *CALB2*, calretinin, *NOS1*, neuronal nitric oxide synthase, *RELN*, reelin, *TAC1, TAC3* and *TAC4*, the tachykinin precursor genes, *GABRA1-5*, GABA_A_R receptor subunits α1-5, *GABRB1-3*, GABA_A_R receptor subunits β1-3, *GABRG1-3*, GABA_A_R receptor subunits γ1-3, *GABRE*, GABA_A_R receptor subunit ε and *GABRD*, GABA_A_R receptor subunit δ

While we used WGCNA to identify clusters of co-expressed genes, we sought to complement those findings with a pairwise correlation analysis. This served both as a validation step and as a method to investigate co-expression patterns between genes of interest that might not pertain to a discrete co-expression cluster. Hence, we performed bivariate correlation analysis of the genes of interest with the *corrplot* package in R 4.0.3 (Figure 1B). This revealed strong correlations (Pearson’s *r* > 0.5, p < 0.05) between *SST* and *GABRA5, GABRA2* and *GABRB1* (Figure 1C-E); between *VIP* and *CCK* (Figure 1F); between *GABRA1* and *GABRB2, GABRG2* and *GABRG3* (Figure 1H-J); and between *PVALB* and genes encoding the subunits of the main GABAergic receptor in the brain, GABA_A_Rα1β2γ2 (*GABRA1, GABRB2* and *GABRG2*) (19)(Figure 1K-M).

### [^11^C]Ro15-4513 and [^11^C]flumazenil PET binding track specific GABAergic microcircuits

Having identified patterns of co-expression between main interneuron markers and GABA_A_R subunits, we then investigated relationships between gene expression and 1) [^11^C]Ro15-4513 (n=10 healthy volunteers) and 2) [^11^C]flumazenil binding (n=16 healthy volunteers). For this purpose, we relied on partial least square (PLS) regression, accounting for spatial autocorrelation. For each radiotracer, we performed two complementary analyses. First, we used as predictors the eigengenes of each of the clusters we identified in the WGCNA analysis. Eigengenes in this context refer to the first principal component of a given cluster, thus representing the pattern of regional expression of all genes within that cluster. Second, we used as predictors all 15,633 genes that passed our pre-processing criteria and inspected the rank of each of our genes of interest in the ranked list of genes according to the spatial alignment of each gene with the tracer. This would provide a sense of how specific the correlation of each of our genes of interest might be as compared to other non-hypothesized genes and the cluster-wise analysis.

### [^11^C]Ro15-4513 binding is associated with *SST*, *GABRA5, GABRA2* and *GABRB1* expression

For [^11^C]Ro15-4513, the first PLS component (PLS1) of the cluster-wise analysis explained alone the largest amount (58.28%, p_spatial_ < 0.0001) of variance in radiotracer binding (Figure S4A). We focused our subsequent analyses on this first component, as it explained the most of variance. This cluster contained *GABRA5, GABRA2, SST* and *GABRB1*, and was assigned the highest positive PLS1 weight (Z=6.18, FDR=1.67 x 10^-8^). In the gene-wise PLS analysis, the first PLS component explained alone the largest amount of variance (57.78%, p_spatial_ < 0.0001) (Figure S4B). *GABRA5*, *GABRB3*, *GABRA2*, *NPY*, *VIP, SST, GABRB1, TAC3, CCK, GABRA3, RELN* and *TAC1* (Z = 4.38-2.28, pFDR = 0.000273-0.0366) were all assigned significant positive weights in descending order (Table 1; for full PLS regression analysis results, see Supplementary table 1). Interestingly, *PVALB* expression had a significant negative PLS1 weight (Z = −2.46, pFDR = 0.0255), which suggested an anti-correlation between *PVALB* and [^11^C]Ro15-4513 binding.

**Table 1.** Weights and significance of covariance between the expression of individual genes of interest and [^11^C]Ro15-4513 signal.

| Gene | PLS rank / 15,633 | PLS gene weight (Z-score) | pFDR |
| --- | --- | --- | --- |
| GABRA5 | 183 | 4.38 | 2.73 x 10<sup>-4</sup> |
| GABRB3 | 275 | 4.20 | 4.11 x 10<sup>-4</sup> |
| GABRA2 | 306 | 4.15 | 4.65 x 10<sup>-4</sup> |
| NPY | 523 | 3.83 | 1.02 x 10<sup>-3</sup> |
| VIP | 527 | 3.82 | 1.02 x 10<sup>-3</sup> |
| SST | 573 | 3.76 | 1.19 x 10<sup>-3</sup> |
| GABRB1 | 804 | 3.48 | 2.41 x 10<sup>-3</sup> |
| TAC3 | 954 | 3.31 | 3.69 x 10<sup>-3</sup> |
| CCK | 1492 | 2.84 | 0.0114 |
| GABRA3 | 1665 | 2.71 | 0.0153 |
| RELN | 2066 | 2.43 | 0.0271 |
| TAC1 | 2283 | 2.28 | 0.0366 |
| PVALB | 13433 | -2.46 | 0.0255 |

Statistically significant results (pFDR<0.05) shown only. PLS weight and pFDR shown to third significant figure. PLS, partial least squares regression analysis.

The radiotracer binding and the distribution of weights resulting from both PLS analyses (cluster-wise and gene-wise) followed and antero-posterior distribution gradient in the brain (Figure 2), consistent with the analogous gradient of *SST* expression shown previously (11). We then followed up these results with a cell-type enrichment analysis, accounting for the weights associated with each gene included in the analysis. This analysis revealed enrichment in genes expressed in SST, CCK and VIP/CCK interneurons (for full results, see Supplement). The result supported an association between the distribution of these cell-types with [^11^C]Ro15-4513 binding.

**Figure 2.**
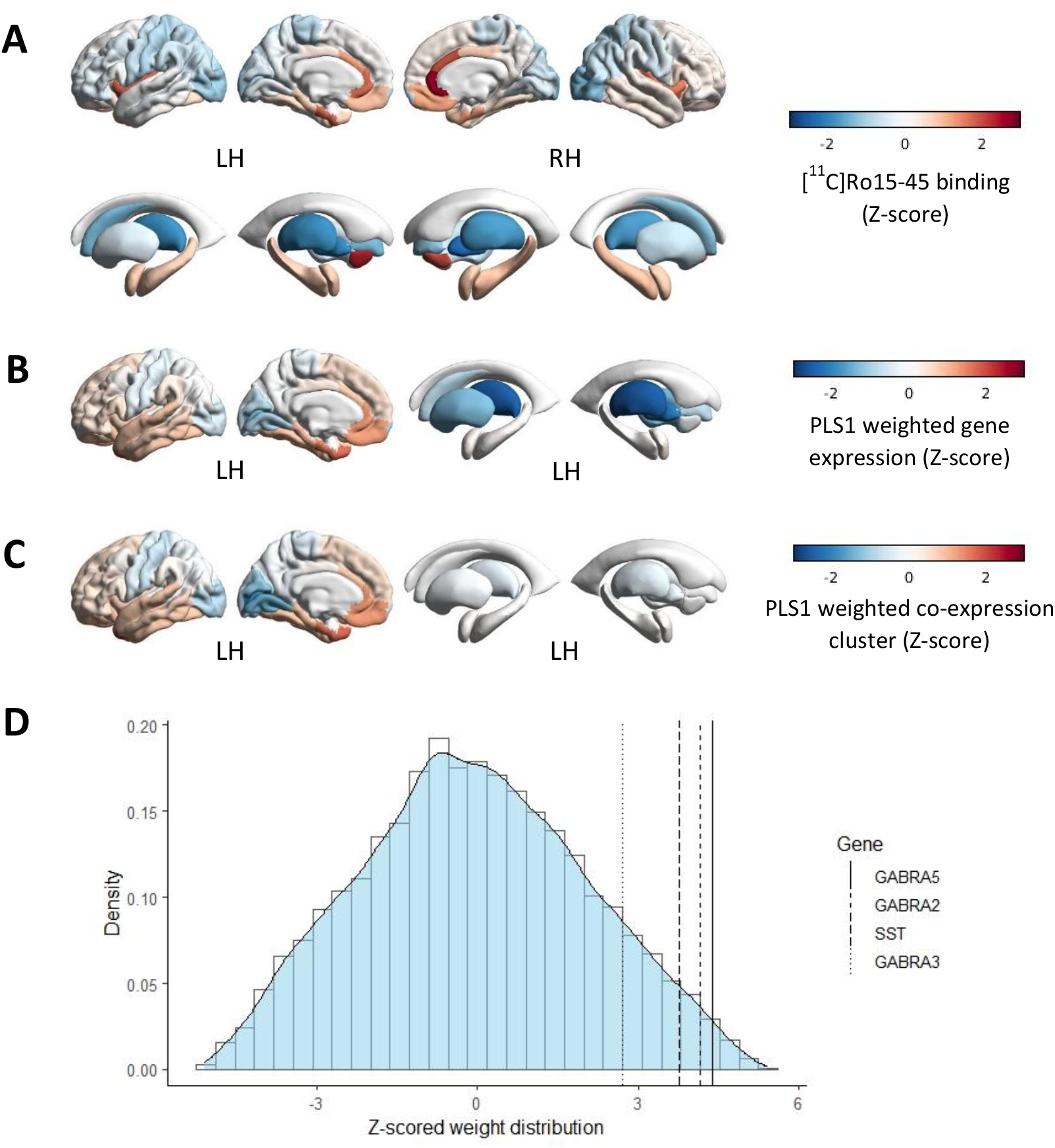
[^11^C]Ro15-4513 binding follows an antero-posterior gradient and spatially tracks *SST, GABRA5, GABRA2* and *GABRA3* expression. Z-scored regional brain distribution of (**A**) [^11^C]Ro15-4513 binding, (**B**) weights of covariance between [^11^C]Ro15-4513 signal and expression of 15,633 genes from the AHBA and (**C**) weights of covariance between [^11^C]Ro15-4513 signal and 52 co-expression clusters from the AHBA. (**D**) density plot of Z-scored weight distribution of 15,633 genes from the AHBA in their covariance with [^11^C]Ro15-4513 signal, with location of *GABRA5, GABRA2, GABRA3* and *SST*. *GABRA2*, GABA_A_R receptor subunit α2, *GABRA3*, GABA_A_R receptor subunit α3, *GABRA5*, GABA_A_R receptor subunit α5, *SST*, somatostatin

### [^11^C]flumazenil binding is associated with *PVALB, GABRA1, GABRB2, GABRG2, GABRG3* and *GAD1* expression

For [^11^C]flumazenil, the first PLS component (PLS1) of the cluster-wise analysis explained alone the largest amount of variance (36.49%, p_spatial_ = 0.001) in radiotracer binding (Figure S4C). This cluster contained *GABRB2, GABRG3, GABRA1, GABRG2* and *GAD1* (Z=7.48, pFDR=1.87 x 10^-12^). In the gene-wise PLS analysis, the first PLS component explained alone the largest amount of variance (37.13%, p_spatial_ = 0.005) (Figure S4D). *GABRB2, GABRD, GABRG3, GABRA1, GABRG2, GABRA4, GAD1, VIP, CCK* and *PVALB* (Z = 6.91-2.26, pFDR = 1.45 x 10^-9^-0.0315) were all assigned significant positive weights in descending order (Table 2; for full PLS results, see Supplementary table 3).

**Table 2.** Weights and significance of covariance between the expression of individual genes of interest and [^11^C]flumazenil signal.

| Gene | PLS rank / 15,633 | PLS gene weight (Z-score) | pFDR |
| --- | --- | --- | --- |
| GABRB2 | 22 | 6.91 | 1.45 x 10<sup>-9</sup> |
| GABRD | 227 | 5.64 | 3.95 x 10<sup>-7</sup> |
| GABRG3 | 392 | 5.11 | 3.97 x 10<sup>-6</sup> |
| GABRA1 | 401 | 5.07 | 4.67 x 10<sup>-6</sup> |
| GABRG2 | 597 | 4.67 | 2.31 x 10<sup>-5</sup> |
| GABRA4 | 2101 | 2.94 | 6.43 x 10<sup>-3</sup> |
| GAD1 | 2405 | 2.70 | 0.0117 |
| VIP | 2566 | 2.60 | 0.0150 |
| CCK | 3050 | 2.26 | 0.0311 |
| PVALB | 3059 | 2.26 | 0.0315 |
| GABRA3 | 12482 | -2.09 | 0.0436 |
| GABRG1 | 12816 | -2.29 | 0.0293 |
| CALB1 | 13166 | -2.50 | 0.0190 |
| NOS1 | 13727 | -2.91 | 6.70 x 10<sup>-3</sup> |
| CALB2 | 15393 | -5.09 | 4.34 x 10<sup>-6</sup> |

Statistically significant results (pFDR<0.05) shown only. PLS weight and pFDR shown to third significant figure. PLS, partial least squares regression analysis.

Radiotracer binding, as well as the distribution of weights resulting from both PLS analyses, followed and postero-anterior distribution gradient in the brain (Figure 3), consistent with analogous pattern of *PVALB* expression shown previously (11). Following up these results with a cell-type enrichment analysis, accounting for weights associated with each gene input into the analysis, revealed enrichment in genes expressed in PVALB, CCK, VIP/CCK and SST interneurons (for full results, see Supplement). The result supported an association between the distribution of PVALB, CCK and VIP/CCK cell-types with [^11^C]flumazenil binding, and suggested an association between genes enriched in the SST cell-type and radiotracer signal despite no direct covariance with *SST* expression found in the PLS analysis.

**Figure 3:**
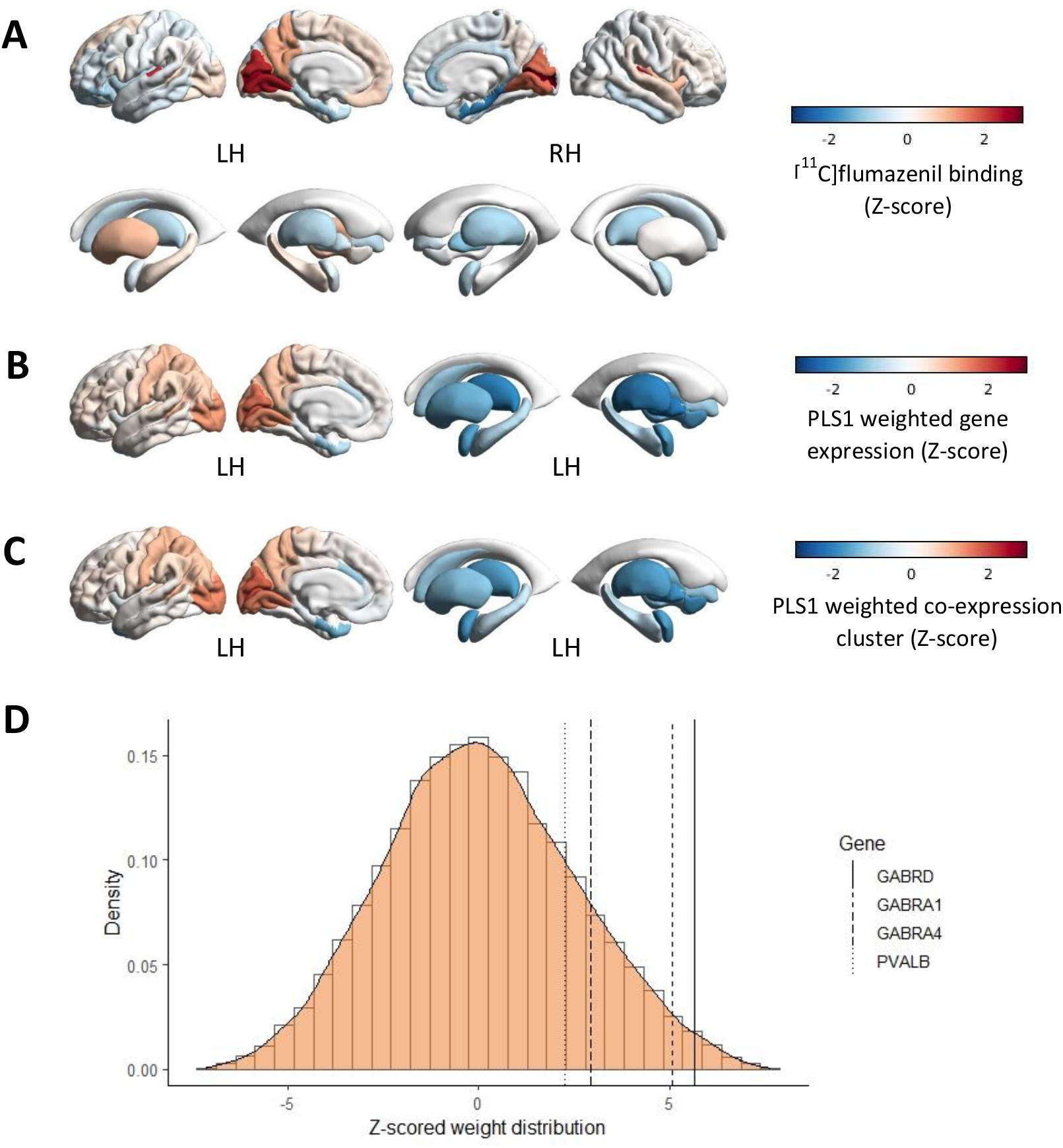
[^11^C]flumazenil binding follows a postero-anterior gradient and spatially tracks *PVALB, GABRD, GABRA1* and *GABRA4* expression. Z-scored brain distribution of (**A**) [^11^C]flumazenil binding, (**B**) weights of covariance between [^11^C]flumazenil signal and expression of 15,633 genes from the AHBA and (**C**) weights of covariance between [^11^C]flumazenil signal and 52 co-expression clusters from the AHBA. (**D**) density plot of Z-scored weight distribution of 15,633 genes from the AHBA in their covariance with [^11^C]flumazenil signal, with location of *GABRD, GABRA1, GABRA4* and *PVALB. GABRA1*, GABA_A_R receptor subunit α1, *GABRA4*, GABA_A_R receptor subunit α4, *GABRD*, GABA_A_R receptor subunit δ, *PVALB*, parvalbumin.

## Discussion

Integrating transcriptional and molecular neuroimaging data in humans, we demonstrate that the spatial pattern of expression of specific GABAergic interneuron markers covaries with that of different GABA_A_R subunits, and that these co-expression patterns explain a substantial portion of the variation in GABA PET radiotracer binding, a measure of *in vivo* brain neurotransmission. Our main finding is that [^11^C]Ro15-4513 and [^11^C]flumazenil signal were differentially associated with the expression of distinct GABAergic interneuron markers and GABA_A_R subunits. While [^11^C]Ro15-4513 followed an anterior distribution that tracked the spatial expression of GABA_A_Rα5 and *SST*, [^11^C]flumazenil followed a more posterior distribution which covaried with the expression of GABA_A_Rα1 and *PVALB*. Overall, these findings have important implications for the study of GABAergic microcircuit dysfunction in psychiatric conditions.

Both GABAergic interneuron distribution and [^11^C]Ro15-4513 and [^11^C]flumazenil binding are spatially heterogeneous across the brain, but the relationship between these micro- and macroscopic processes remained poorly understood. Previous PET studies described that [^11^C]Ro15-4513 and [^11^C]flumazenil binding were anti-correlated along an anterior-posterior axis (23), resembling a largely developmentally preserved gradient of *SST* to *PVALB* distribution in the human brain, as recently reported (11). Using cutting-edge imaging transcriptomics, we show that these findings may not be coincidental, and that specific GABAergic microcircuits may be investigated in humans *in vivo* with existing neuroimaging methods.

Our approach relies on indirect spatial associations between PET radiotracer binding and gene expression across brain regions. Hence, direct extrapolations about specific synapse contribution to our findings should be made with caution. Nevertheless, our results are corroborated by existing preclinical research using more precise molecular methods. For instance, we observed that the spatial pattern of [^11^C]Ro15-4513 binding covaried most strongly with a gene cluster containing *GABRA5, GABRA2* and *SST*. This finding is consistent with preclinical literature showing that GABA_A_Rα5 are enriched on principal cell membranes targeted by SST cells (5,26,40). Moreover, [^11^C]Ro15-4513 has been shown to present 10-15-fold higher affinity to human cloned GABA_A_Rα5 than to GABA_A_Rα1-3 (23) and the co-expression of *GABRA2* and *GABRA5* was previously shown by immunohistochemistry in the rat brain (16), which we observed in our analyses as well. Interestingly, we also found covariance between the expression of *CCK* and *GABRA3* and the distribution of [^11^C]Ro15-4513 signal. GABA_A_Rα2/3 expression in rodents has been most consistently found in the post-synapse of principal cells targeting CCK basket cells (3,19,24,41). It is plausible that the association between [^11^C]Ro15-4513 binding and CCK cell-related microcircuits we found is circumstantial, if enough spatial overlap between the two microcircuits exists. Alternatively, the discrepancy could be a result of secondary affinity of the tracer to GABA_A_Rα2/3, as the latter constitute less than 5% of all GABA_A_Rs in the brain (41) and PET radiotracers are administered systemically with a bolus injection. Future research using precise methods such as immunocytochemistry or autoradiography with pharmacological blocking is warranted to determine whether [^11^C]Ro15-4513 binds to GABA_A_Rα2/3 in addition to GABA_A_Rα5.

The spatial pattern of [^11^C]flumazenil binding covaried most strongly with the cluster containing *GAD1, GABRA1, GABRB2, GABRG2* and *GABRG3*. Prior evidence that [^18^]flumazenil accumulation across the mouse brain after mutations in α2, α3 and α5 subunits but not in α1 remained similar to that in wild-type mice suggests that flumazenil binding to GABA_A_Rα1 predominates in the mammalian brain (42). Our finding is also in agreement with the notion that the abundance of GABA_A_Rα1 in the brain may be reflected in greater GABA_A_Rα1 flumazenil binding (14,23). Indeed, GABA_A_Rα_1_β_2_γ_2_ is the most widely expressed GABA_A_R in brain (8,20) and the co-expression of *GABRA1, GABRB2* and *GABRG2* is supported by analogous observation in preclinical immunohistochemistry studies (43,44). Interestingly, [^11^C]flumazenil signal was also associated with *PVALB* expression, consistent with our observation of a high association between *PVALB* expression with *GABRA1*, *GABRB2* and *GABRG2*, and in line with preclinical evidence that GABA_A_Rα1 is located in synapses formed by PVALB interneurons onto each other and onto principal cells (19,24–26,41). Finally, covariance of both *GABRA4* and *GABRD* expression with [^11^C]flumazenil binding is consistent with the finding that those subunits are commonly co-expressed in the forebrain and that the δ subunit associates extrasynaptically with α1, primarily in the cerebellum but also in the cerebrum (45,46) where [^11^C]flumazenil uptake is higher than that of [^11^C]Ro15-4513 (23).

Other noteworthy patterns of gene expression covariance and cell-type enrichment with radiotracer signal patterns emerged. For instance, the lack of *PVALB* assignment to a cluster containing any other genes of interest may reflect the widespread expression of this protein in the human brain. Additionally, the binding of both radiotracers was associated with *CCK* and *VIP* expression. The latter observation may reflect a co-localisation of several cell types within regions showing radiotracer binding, since VIP interneurons primarily innervate SST interneurons (3,26) and VIP/CCK cells were shown to target principal cells and PVALB basket cells (47). On the other hand, the *VIP* co-expression cluster was enriched in excitatory cell type markers. These observations warrant further investigation with more precise methods such as immunocytochemistry, chemogenetics or pharmacological manipulations.

Our findings have important implications for future studies investigating GABAergic dysfunction with PET. Abnormalities in both PVALB interneuron/GABA_A_Rα1 and SST interneuron/GABA_A_Rα2,3,5 pathways have been hypothesised to contribute to the pathophysiology of several brain disorders, including those with affective abnormalities (8). For instance, GABA_A_Rα2 and GABA_A_Rα3 agonism is implicated in benzodiazepine-mediated anxiolysis (14,19). To our knowledge there are no existing studies in anxiety disorders using [^11^C]Ro15-4513, which we found to track the spatial pattern of GABA_A_Rα2 and GABA_A_Rα3 expression. This suggests that GABA_A_Rα2 and GABA_A_Rα3 deficits may be involved in anxiety disorders, which could be investigated with [^11^C]Ro15-4513. Such studies would complement existing GABA PET studies in anxiety disorders, which used [^11^C]flumazenil to detect decreased binding in the insula in panic disorder and other anxiety disorders (48–50). As we found [^11^C]flumazenil to correlate with GABA_A_Rα1 expression specifically, and this receptor is associated with undesired sedative effects of benzodiazepines (14,19), [^11^C]Ro15-4513 may be used in drug discovery paradigms for new anxiolytic medications with more GABA_A_Rα2/3 selectivity (51). SST interneuron dysfunction has also been implicated in the aetiology of depressive disorders (41,52), and a preclinical model of SST cell disinhibition produced an anxiolytic- and antidepressant-like effect akin to that of benzodiazepines or ketamine (53). As α5 subunit increases have been found *post-mortem* in patients with depression (9), [^11^C]Ro15-4513 studies may also inform SST interneuron/GABA_A_Rα5 microcircuit dysfunction in this condition.

Furthermore, our observation that [^11^C]Ro15-4513 and [^11^C]flumazenil track SST and PVALB interneuron-related microcircuits, respectively, informs research in brain disorders in which hippocampal dysfunction is thought to play a key role, such as schizophrenia (7,10). The hippocampus is one of the few regions where *SST* and *PVALB* are not anti-correlated (11), as is enriched in both microcircuits (10). SST and PVALB interneuron loss were reported in hippocampi of patients with schizophrenia by *post-mortem* examination (54), implicating both interneuron types in the observed hyperactivity of this region in patients *in vivo* (10). However, [^11^C]flumazenil PET studies in schizophrenia have produced inconsistent results across multiple cortical regions and not the hippocampus (55). Conversely, a recent [^11^C]Ro15-4513 study in schizophrenia found decreases in radiotracer binding in the hippocampus of antipsychotic medication-free patients (56). Analogous deficits were not identified in currently medicated patients (56,57), consistent with the observation that antipsychotic treatment increases [^11^C]Ro15-4513 binding specifically in the rat hippocampus (58). Interestingly, people at clinical high-risk for psychosis show increased hippocampal perfusion (59) which is correlated with prefrontal GABA levels, particularly in those individuals who subsequently developed psychosis (60). Taken together, these findings may suggest that SST interneuron dysfunction in the hippocampus and PVALB interneuron abnormalities in cortical regions may play a role in the onset of psychosis. Future PET studies in early psychosis may address this hypothesis by examining PVALB interneuron-mediated networks with [^11^C]flumazenil, and SST interneuron-associated inhibition with [^11^C]Ro15-4513.

Our study has some limitations worth acknowledging. First, we relied on indirect spatial associations between gene expression and radiotracer binding, which alone does not directly imply co-expression in the same cell or in interconnected neurons, nor direct radiotracer binding. However, we note that our findings are broadly supported by the preclinical literature using more fine-grained methods, which lends support to the plausibility of our imaging transcriptomics findings in humans. Our findings were generated though several hypotheses, which may guide focused molecular studies in the *post-mortem* human brain, and which can be easily extrapolated to other neurochemical systems. Moreover, our work follows a similar approach to previous studies investigating relationships between interneuron marker expression and resting-state activity, (11) and benzodiazepine receptor availability (61). Second, the AHBA includes data from six donors only. Samples from the right hemisphere were only collected for two donors, which led us to restrict our analyses to the left hemisphere. Although not a specific limitation of this study, this raises questions about whether this small sample can capture well the principles of organisation of the canonical architecture of gene expression in the human brain and generalise well. Finally, because we applied an intensity threshold to the microarray dataset to minimise inclusion of unreliable measures of gene expression, we were not able to investigate some genes of interest, including *GABRA6, GABRP, GABRQ* and *GABRR1-3*, due to the low intensity of signal for these genes in the AHBA dataset as compared to background. Future studies using high sensitivity methods to measures expression of these genes across the whole brain will help to complement our findings in this respect.

In summary, we provide evidence of spatial alignment between the expression of: 1) *SST*, *GABRA5* and *GABRA2;* 2) *PVALB* and *GABRA1, GABRB2* and *GABRG2;* and 3) *VIP* and *CCK* in human. These findings expand our understanding of the canonical transcriptomic architecture of different GABAergic interneuron microcircuits in the human brain. Furthermore, we provide first evidence that these separate interneuron subtype-specific microcircuits covary with [^11^C]Ro15-4513 and [^11^C]flumazenil binding in a largely non-overlapping manner. While [^11^C]Ro15-4513 signal covaried with the regular-spiking interneuron marker *SST* and genes encoding several major benzodiazepine-sensitive GABA_A_R subunits implicated in affective functioning (*GABRA5, GABRA2* and *GABRA3*), [^11^C]flumazenil tracked the fast-spiking interneuron marker *PVALB* and genes encoding subunits comprising the most widely expressed receptor (GABA_A_Rα1β2γ2), linked to general neuronal network activity. These findings have important implications for existing and future PET studies of GABA dysfunction in psychiatric and neurological disorders, as they may inform methodological choices for imaging the GABAergic system, and help the interpretation of findings within a framework that bridges the gap between genes, cells and macroscopic molecular neuroimaging features *in vivo*.

## Methods

### The Allen Human Brain Atlas (AHBA) dataset

The AHBA dataset includes microarray data of gene expression in *post-mortem* brain samples from six healthy donors (one female, mean age +/− SD 42.5 +/− 13.38, range 24-57) (36). The brain (cerebrum including the brainstem) was sampled systematically across the left hemisphere in all six donors, and the right hemisphere in two of the donors. Manual macrodissection was performed on the cerebral and cerebellar cortex, as well as subcortical nuclei, in 50-200mg increments. Subcortical areas and cerebellar nuclei were sampled with laser microdissection in 36mm^2^ increments. RNA was then isolated from these dissections and gene expression quantified with microarray. More information about donor characteristics and dataset generation can be found in the Allen Human Brain Atlas website (https://human.brain-map.org/).

### Gene expression data: preprocessing and spatial mapping

Human gene expression microarray data was extracted from the AHBA with the *abagen* toolbox (https://github.com/netneurolab/abagen) (62) in JupyterLab Notebook through anaconda3 in Python 3.8.5. We mapped AHBA samples to the parcels of the Desikan-Killiany, including 83 brain regions across both brain hemispheres (34 cortical and 7 subcortical regions per brain hemisphere, plus brainstem). Genetic probes were reannotated using information provided by Arnatkeviciute et al., 2019 (63) instead of the default probe information from the AHBA dataset to exclude probes that cannot be reliably matched to genes. According to the existing guidelines for probe-to-gene mappings and intensity-based filtering (63), the reannotated probes were filtered based on their intensity relative to background noise level; probes with intensity less than background in ≥50% of samples were discarded. A single probe with the highest differential stability, ΔS(p), was selected for each gene, where differential stability was calculated as (64):

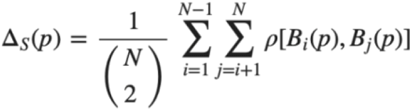
where ρ is Spearman’s rank correlation of the expression of a single probe p across regions in two donor brains, Bi and Bj, and N is the total number of donor brains. This procedure retained 15,633 probes, each representing a unique gene.

Next, tissue samples were assigned to brain regions using their corrected MNI coordinates (https://github.com/chrisfilo/alleninf) by finding the nearest region within a radius of 2 mm. To reduce the potential for misassignment, sample-to-region matching was constrained by hemisphere and cortical/subcortical divisions. If a brain region was not assigned to any sample based on the above procedure, the sample closest to the centroid of that region was selected in order to ensure that all brain regions were assigned a value. Samples assigned to the same brain region were averaged separately for each donor. Gene expression values were then normalized separately for each donor across regions using a robust sigmoid function and rescaled to the unit interval. Scaled expression profiles were finally averaged across donors, resulting in a single matrix with rows corresponding to brain regions and columns corresponding to the retained 15,633 genes.

The genes of interest list included data on all available GABA_A_R subunits and interneuron markers defined according to the Petilla terminology (13) and the existing animal literature. These were: the GABA-synthesising enzymes GAD67 (*GAD1*) and GAD65 (*GAD2*) (29), parvalbumin (*PVALB*) (25), somatostatin (*SST*) (30), vasoactive intestinal peptide (*VIP*) (31), cholecystokinin (*CCK*), neuropeptide Y (*NPY*) (32), calbindin (*CALB1*) (30), calretinin (*CALB2*) (30), neuronal nitric oxide synthase (*NOS1*) (3), reelin (*RELN*) (33), and the tachykinin precursor genes *TAC1, TAC3* and *TAC4* (34). The genes of interest that did not pass this intensity-based thresholding were *GABRA6, GABRP, GABRQ* and *GABRR1-3*.

### Weighted gene co-expression network analysis (WGCNA)

Hierarchical clustering of genes by their expression across brain regions was performed with the WGCNA package (35) in R 4.0.3 for each gene expression dataset. The ‘signed’ WGCNA method was chosen to form clusters (modules) enriched in genes which expression was positively correlated, indicating co-expression (35). Gene expression correlation matrix was transformed into an adjacency matrix using the soft threshold power of 14. This power value was chosen as it was the first value at which the network satisfied the free-scale topology criterion at R^2^ > 0.8, therefore maximising mean network node connectivity (see Supplementary Figure S5). The adjacency matrix was then transformed into a dissimilarity measure matrix, representing both the expression correlation between pairs of genes as well as the number of the genes they both highly correlated with positively (‘neighbours’) (35). Finally, average-linkage hierarchical clustering using the dissimilarity measure was performed. Individual modules were identified through the classic ‘tree’ dendrogram branch cut (65).

### Bivariate correlation of genes of interest

To investigate correlations between individual pairs of genes of interest, bivariate correlation analysis was performed and visualised in R 4.0.3 using the *Hmisc* and *corrplot* packages. All available genes of interest were input into a bivariate correlation analysis. The p-value threshold was set to p<0.05.

### Parametric map of [^11^C]Ro15-4513 binding

Ten healthy participants (four females, mean age +/− SD 25.40 +/− 3.20, range 22-30) with no history of psychiatric diagnoses, neurological illness or head trauma with loss of consciousness were scanned with the radiotracer [^11^C]Ro15-4513. Scanning was performed on a Signa tm PET-MR General Electric (3T) scanner at Invicro, A Konica Minolta Company, Imperial College London, UK. The study was approved by the London/Surrey Research Ethics Committee. All subjects provided written informed consent before participation, in accordance with The Declaration of Helsinki. The radiotracer was administered through the dominant antecubital fossa vein in a single bolus injection, administered at the beginning of the scanning session. The maximum amount of radiation administered was 450MBq. PET acquisition was performed in 3D list mode for 70 minutes and binned in the following frames ADD. Attenuation correction was performed with a ZTE sequence (voxel size: 2.4×2.4×2.4mm^3^, field of view=26.4, 116 slices, TR=400ms, TE=0.016ms, flip angle=0.8°). A T1-weighted IR-FSPGR sequence was used for PET image co-registration (voxel size: 1×1×1 mm^3^, field of view=25.6, 200 slices, TR=6.992ms, TE=2.996ms, TI=400ms, flip angle=11°).

Individual subject images were generated with MIAKAT v3413 in Matlab R2017a. For each subject, an isotropic, skull-stripped IR-FSPGR structural image normalised to the MNI template was co-registered onto an isotropic, motion-corrected integral image created from the PET time series. Binding potential parametric maps were estimated through a simplified reference tissue model using the pons as the reference region and solved with basis function method (66). The individual parametric maps were averaged using SPM imCalc function and resliced with the Co-register: Reslice function to match the dimensions of the Desikan-Killiany atlas (voxel size 1×1×1mm, number of voxels per direction X=146, Y=182, Z=155). Finally, the averaged parametric map of [^11^C]Ro15-4513 binding was resampled into 83 regions of the Desikan-Killiany atlas space using the *fslmeants* function from FSL.

### Parametric map of flumazenil binding

An averaged parametric map of maximal binding of the [^11^C]flumazenil ([^11^C]Ro15-1788) radioligand was downloaded from an open-access dataset made available by the Neurobiology Research Unit at Copenhagen University Hospital (https://xtra.nru.dk/BZR-atlas/). In brief, 16 healthy participants between 16-46 years old (nine females, mean age +/− SD 26.6 +/− 8) were scanned on a CTI/Siemens High-Resolution Research Tomograph. Regional radiotracer binding was estimated using Logan analysis. For full details on the generation of this map, please refer to the original publication (61).

### Covariance between [^11^C]Ro15-4513 and [^11^C]flumazenil PET radiotracer binding and gene expression

Partial least squares regression (PLS) analysis was used to identify genes whose expression was most strongly associated with either [^11^C]Ro15-4513 or [^11^C]flumazenil binding. The script used for this analysis is available elsewhere (67) and was run using Matlab R2017a. The predictor variable matrix comprised gene expression per brain region in the left hemisphere only since the AHBA only includes data from the right hemisphere for two out of the six donors. The response variable matrices comprised [^11^C]Ro15-4513 and [^11^C]flumazenil binding, respectively, in the 42 brain regions of the left hemisphere. The analysis was then repeated using the 52 WGCNA module eigengenes as the predictor variables. Prior to each PLS analysis, both predictor and response matrices were Z-scored.

The first PLS component (PLS_1_) is the linear combination of the weighted gene expression scores that have a brain expression map that covaries the most with the map of tracer binding. As the components are calculated to explain the maximum covariance between the dependent and independent variables, the first component does not necessarily need to explain the maximum variance in the dependent variable. However, as the number of components calculated increases, they progressively tend to explain less variance in the dependent variable. Here, we tested across a range of components (between 1 and 15) and quantified the relative variance explained by each component. The statistical significance of the variance explained by each component was tested by permuting the response variables 1,000 times, while accounting for spatial autocorrelation using a combination of spin rotations for the cortical parcels and random shuffling for the subcortical ones. We decided to focus on the component explaining the largest amount of variance, which in our case was always the first component (PLS_1_). The error in estimating each gene’s PLS_1_ weight was assessed by bootstrapping, and the ratio of the weight of each gene to its bootstrap standard error was used to calculate the *Z* scores and, hence, rank the genes according to their contribution to PLS_1_. The code used to implement these analyses can be found in https://github.com/SarahMorgan/Morphometric_Similarity_SZ.

## Acknowledgements

The authors would like to extend special thanks to colleagues from the Neurobiology Research Unit at Copenhagen University Hospital for making their data publicly available, as well as Dr Samuel Cooke for his advice on the interpretation of the results.

## Author contributions

PBL, DM, FET and GM designed the research, PBL conducted the research, DM and MV provided analytic support, PBL wrote the manuscript and made all figures. All authors edited the manuscript and made contributions to the interpretation of the data.

## Code availability

Code used for any part of the project can be made available at request.

