## Supplementary material for "Cellular and molecular signatures of *in vivo* GABAergic neurotransmission in the human brain"

### Supplementary results

#### Cell type specificity analysis on WGCNA co-expression clusters in AHBA

We used the Over-Representation Analysis method in the WEB-based GEne SeT AnaLysis Toolkit (WebGestalt, www.webgestalt.org) (1) to determine cell type enrichment in the clusters identified through our analysis. We focused on the clusters containing three major non-overlapping interneuron markers: *SST*, *PVALB* and *VIP*. The cluster including *SST* from the AHBA-based analysis was enriched in the In4a (CNR1/RELN positive interneurons), In2 (CCK/RELN/CALB2), In1a (CCK/RELN) and In4b (CCK) interneuron subtypes (Figure S1A). The *PVALB* cluster was enriched in the In6b (PVALB/TAC1) cell type (Figure S1B). Finally, the cluster including the markers *VIP* and *CCK* showed enrichment in multiple excitatory cell types (Figure S1C).

**Figure S1.** Cell type enrichment of the co-expression clusters from the Allen Human Brain Atlas (AHBA) for the three main interneuron markers. Enrichment of **A** the somatostatin co-expression cluster; **B** the parvalbumin co-expression cluster; and **C** of the vasoactive intestinal peptide co-expression cluster. Enrichment score indicates the strength of association between lists of genes co-expressed in cell types identified by previous single-cell transcriptomic analysis (2) and individual co-expression clusters returned by our WGCNA analysis.

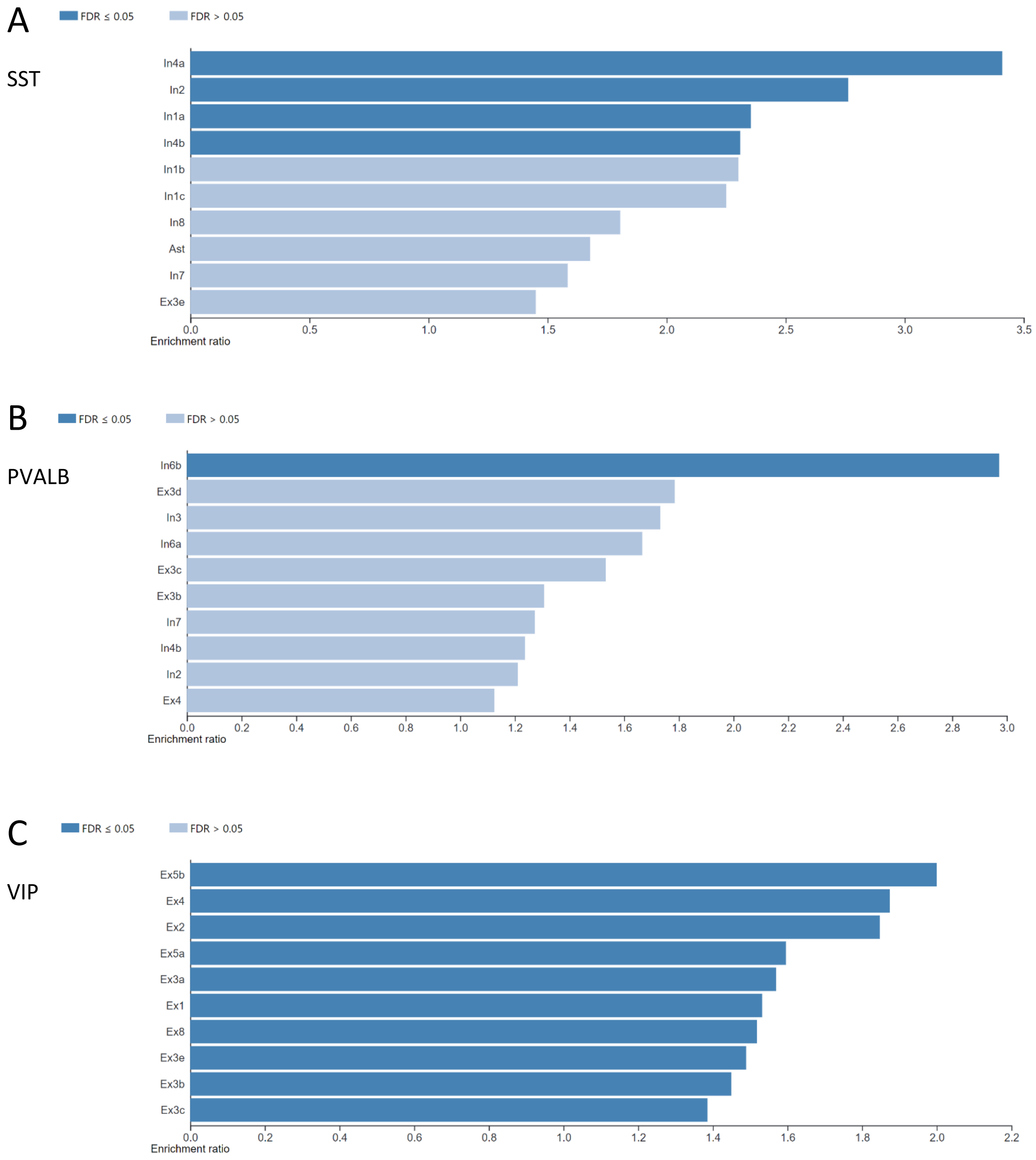

#### Cell type specificity analysis on PLS regression analysis of PET radiotracer maps and gene expression from the AHBA

To further validate our approach and investigate the potential cell type enrichment of the results of our PLS regression analyses, we performed additional analyses (WebGestalt Gene Set Enrichment Analyses) on the two gene-wise PLS analysis results for [$^{11}$C]Ro15-4513 and for [$^{11}$C]flumazenil. [$^{11}$C]Ro15-4513 binding was associated with genes enriched in the interneuron subtypes In8 (SST cells), In1c (CCK/VIP/TAC3/CALB2), In4b (CCK), In1a (CCK/RELN), In1b (CCK) and In3 (negative for SST, PVALB, CCK or VIP) (Figure S2). It was also associated with several excitatory (Ex) cell types but no non-neuronal cell types.

**Figure S2.** Cell type enrichment of the 15,633 genes from the Allen Human Brain Atlas according to their covariance with [$^{11}$C]Ro15-4513 signal. Positive (blue) or negative (orange) normalised enrichment score indicates the strength of association between lists of genes co-expressed in cell types identified by previous single-cell transcriptomic analysis (2) and the full list of genes from the AHBA dataset with their associated weights from the PLS analysis.

End, endothelial cells, Ex3e, Ex4, Ex6b, Ex8, excitatory neuron type 3e, 4, 6b and 8, In1a, In1b, In1c, In3, In4b, In8, interneuron type 1a, 1b, 1c, 3, 4b and 8, Mic, microglia, Oli, oligodendrocytes, Per, pericytes

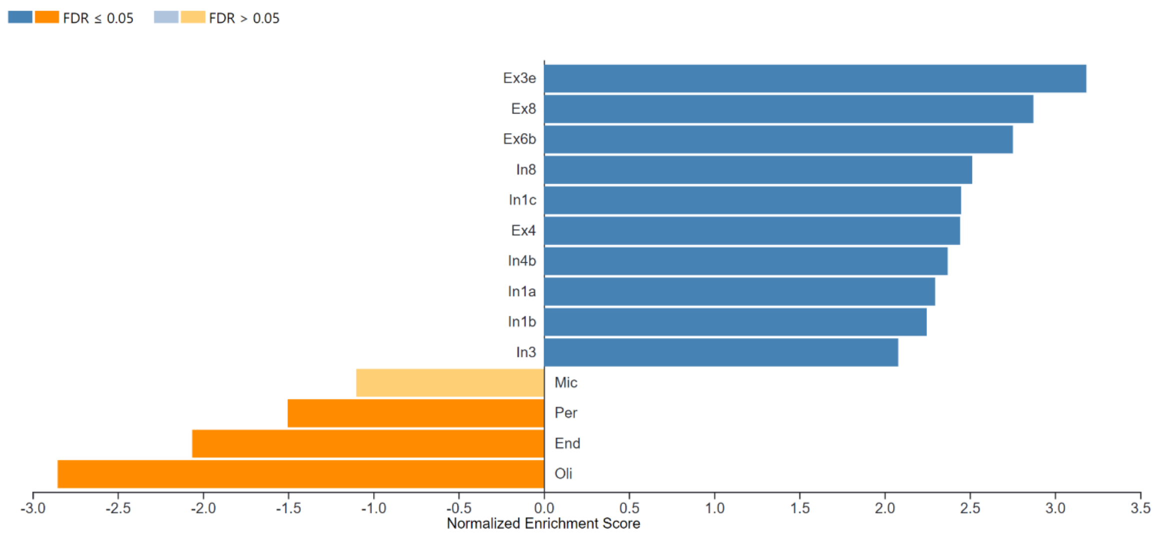

For the corresponding WEBgestalt Gene Set Enrichment Analysis, [$^{11}$C]flumazenil binding was associated with genes enriched in the interneuron subtypes In3 (negative for SST, PVALB, CCK or VIP), In6a (PVALB cells), In4b (CCK), In1c (CCK/VIP/TAC3/CALB2), In8 (SST) and In2 (CCK/RELN/CALB2) (Figure S3). It was also associated with several excitatory (Ex) cell types but no non-neuronal cell types.

**Figure S3.** Cell type enrichment of the 15,633 genes from the Allen Human Brain Atlas according to their covariance with [$^{11}$C]flumazenil signal. Positive (blue) or negative (orange) normalised enrichment score indicates the strength of association between lists of genes co-expressed in cell types identified by previous single-cell transcriptomic analysis (2) and the full list of genes from the AHBA dataset with their associated weights from the PLS analysis.

Ast, astrocytes, End, endothelial cells, Ex3e, Ex3d, Ex6b, Ex8, excitatory neuron type 3e, 3d, 6b and 8, In1c, In2, In3, In4b, In6a, In8, interneuron type 1c, 2, 3, 4b, 6a and 8, Mic, microglia, Oli, oligodendrocytes, OPC, oligodendrocyte precursor cells, Per, pericytes

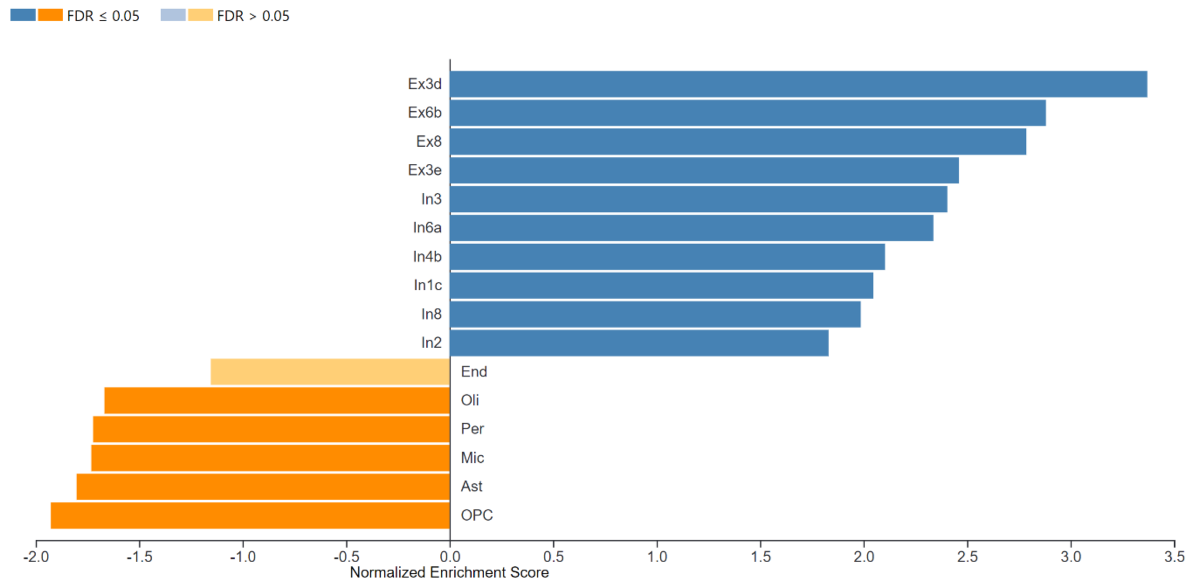

**Figure S4.** Percentage of variance explained by principal components resulting from the partial least squares regression (PLS) analysis for **A** cluster-wise PLS for [$^{11}$C]Ro15-4513, **B** gene-wise PLS for [$^{11}$C]Ro15-4513, **C** cluster-wise PLS for [$^{11}$C]flumazenil, **D** gene-wise PLS for [$^{11}$C]flumazenil.

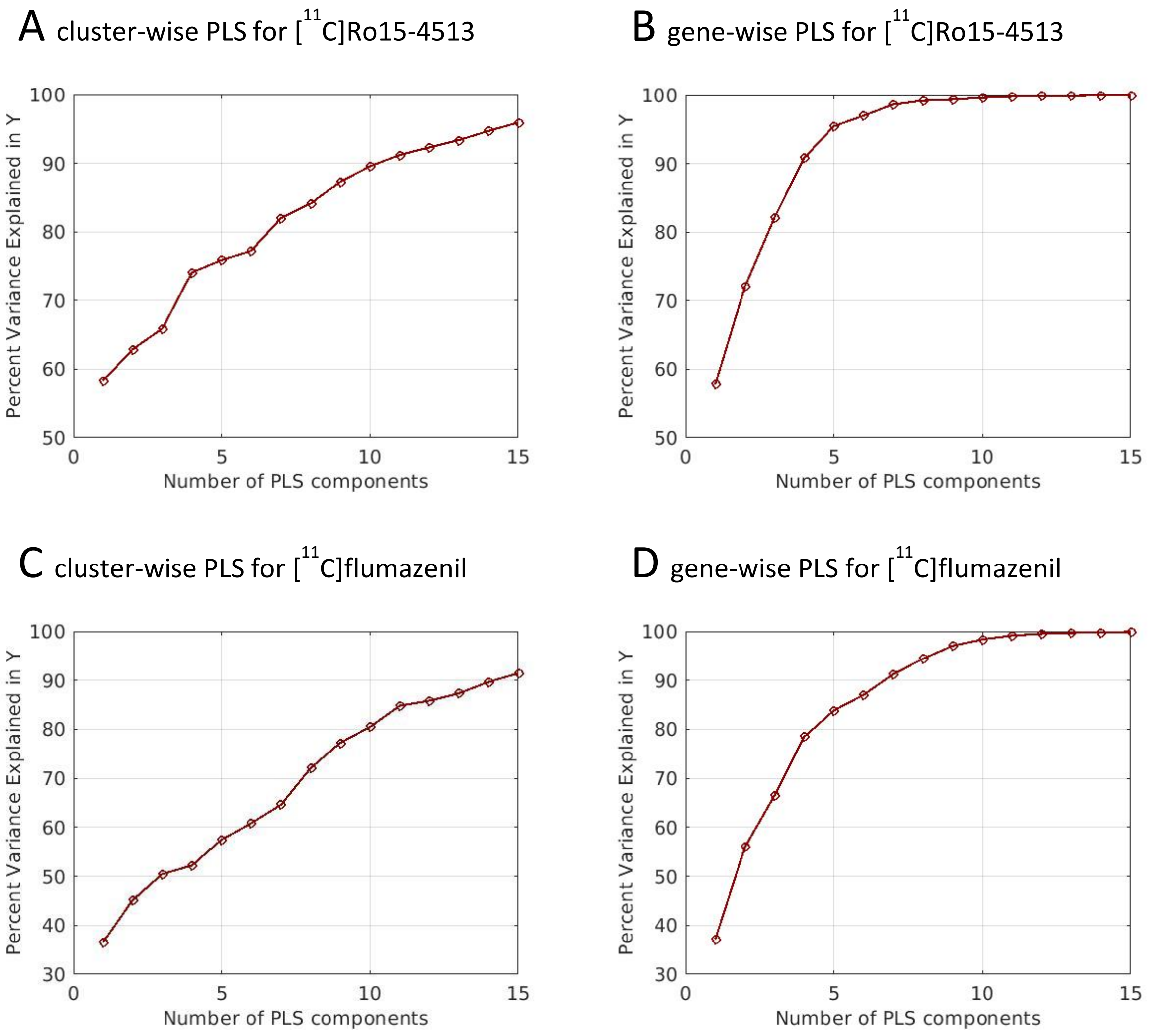

Supplementary Table 1. Results from the first principal component of the gene-wise PLS regression analysis for [$^{11}$C]Ro15-4513.

Supplementary Table 2. Results from the first principal component of the cluster-wise PLS regression analysis for [$^{11}$C]Ro15-4513.

Supplementary Table 3. Results from the first principal component of the gene-wise PLS regression analysis for [$^{11}$C]flumazenil.

Supplementary Table 4. Results from the first principal component of the cluster-wise PLS regression analysis for [$^{11}$C]flumazenil.

### Supplementary methods

#### WGCNA parameter choice

The WGCNA gene expression correlation matrix was transformed into an adjacency matrix using the soft threshold power of 14. This power value was chosen as it was the first value at which the network satisfied the free-scale topology criterion at $R^2 > 0.8$, therefore maximising mean network node connectivity (Figure S5).

**Figure S5.** Power choice function for the weighted gene co-expression network analysis for the Allen Human Brain Atlas database.

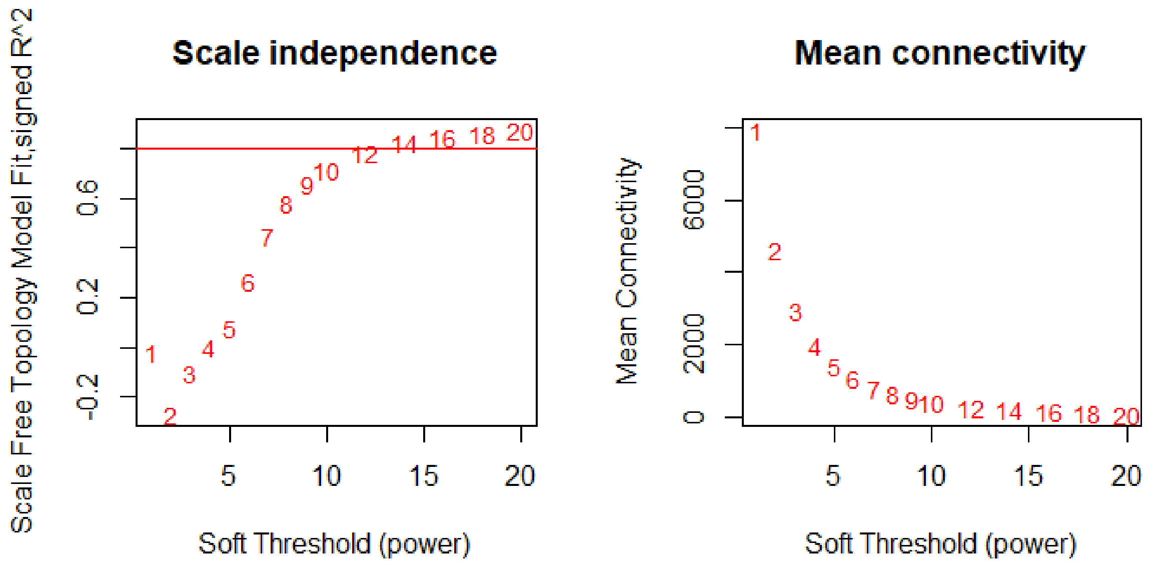


#### Cell type enrichment analyses – WGCNA clusters

Cell type enrichment analysis of the clusters containing main interneuron markers (*SST*, *PVALB* and *VIP*) returned by WGCNA for the AHBA was performed in WEB-based Gene SeT AnaLysis Toolkit (WebGestalt, www.webgestalt.org) (1). The organism of interest was set to Homo sapiens, chosen method was Over-Representation Analysis, and the functional database was derived from brain-cell type single cell gene expression, as identified in a previous study. For each cluster, the gene list extracted from the analysis in R was uploaded into the software. The reference gene set comprised all the 16,533 genes in the AHBA dataset.

#### Cell type enrichment analyses – gene-wise PLS results of radiotracer binding

Cell type enrichment analysis of the two gene-wise PLS analyses for Ro15 and flumazenil were also performed using (WebGestalt, www.webgestalt.org) (1). The Gene Set Enrichment Analysis was chosen to adapt analysis method to input type. We input the complete resulting list of 15,633 genes with their associated weights into a Gene Set Enrichment Analysis in the WEB-based Gene SeT AnaLysis Toolkit We input data on 35 cell types from a previous sequencing experiment by Lake et al (2) as the functional database for cell type assignment.
